# RNA-binding proteins control the G2-M checkpoint of the germinal centre B cell

**DOI:** 10.1101/2024.12.01.626220

**Authors:** Fiamma Salerno, Alex J. Whale, Louise S. Matheson, William S. Foster, Melanie Stammers, Sarah E. Bell, Daniel J. Hodson, Michelle A. Linterman, Jonathan Houseley, Martin Turner

## Abstract

How germinal centre (GC) B cells undergo rapid cell division while maintaining genome stability is poorly understood. Here, we show that the RNA-binding proteins ZFP36L1 and ZFP36L2 act downstream of antigen-sensing and protect GC B cells from replication stress by controlling a cell cycle-related RNA post-transcriptional regulon. ZFP36L1 and ZFP36L2 safeguard faithful completion of mitosis by restraining the expression of CDK1 and cyclin B1, whilst controlling their activity through regulation of a p21-mediated negative feedback loop. In the absence of ZFP36L1 and ZFP36L2, GC B cells arrest in G2-M and die by apoptosis, resulting in curtailed GC responses. This is associated with stalling of the DNA replication fork at active replication initiation zones, which causes replication stress and increased activity of the ATR/CHK1 DNA damage response. Our findings reveal that gene regulation by RNA-binding proteins is essential for a functional G2-M checkpoint to operate in GC B cells.

## Introduction

Germinal centres (GC) are unique microanatomical structures within secondary lymphoid organs where antigen-specific B cells clonally expand and undergo somatic hypermutation (SHM) of their immunoglobulin genes. This mutational process, coupled with selection signals from follicular T helper cells and follicular dendritic cells, pushes GC B cells through multiple rounds of antigen-based selection, clonal expansion and immunoglobulin diversification. Together, these events drive affinity maturation and are essential for the development of effective antibody responses^1, 2^.

Prolific cell division is a characteristic feature of the GC, with about 30% of GC B cells being in cycle at any time^3, 4^. Their fast and intense proliferation is tightly regulated while B cells transit between the light zone (LZ) and the dark zone (DZ) of the GC, and is matched by high rates of cell death^5^ to limit GC size. T cell help leading to MYC and cyclin D3 expression in LZ B cells has been proposed to control the speed of DNA replication and cell cycle phase transitions^4, 6, 7, 8^. The intensity of T cell help received within the LZ determines the number of times selected B cells divide upon transition to the DZ^7, 9^. However, fast dividing DZ B cells have to exit the cell cycle and can linger for a few hours before re-entering the LZ in search for T cell help^3, 4^. Once in the LZ, GC B cells remain in G1 while they wait for selection and progress to the S phase^3, 10, 11^. Proliferation of GC B cells is therefore punctuated by a transient quiescent phase and integrated with pathways of cell death, with DZ B cells in the late G1 phase being particularly prone to apoptosis^3, 4, 10^.

The cell cycle of GC B cells is also connected to SHM, as the most proliferative B cells display a greater number of affinity-enhancing mutations in their immunoglobulin genes^6, 12, 13^. Immunoglobulin diversification is achieved through repair of activation-induced deaminase (AID) initiated mutations by multiple error-prone DNA repair mechanisms^12^. How cell cycle checkpoints are enforced in GC B cells to ensure cell survival is a key part of affinity maturation but remains incompletely understood. The expression of the transcription factor BCL6 is one mechanism to protect B cells from cell cycle checkpoint-mediated death, as it represses the ATR-CHK1-P53 axis^14^. However, deletion or reduced expression of CHK1 in B cells has also been shown to impair GC responses^15, 16^ and to cause accumulation of chromosomal aberrations in activated B cells from *Chk1*^+/−^*Chk*2^+/−^ or *Chk1*^+/−^*Chk2*^−/−^ mice^17^. Understanding how fluctuation between proliferation and quiescence cycles are connected to DNA damage responses is therefore important for immunity and the malignant transformation of GC B cells.

Cell cycle progression requires rapid oscillation in the amounts of many proteins, which is achieved by interconnecting transcription, translation and protein degradation. Because cell cycle proteins have very short half-lives, the output of RNA translation becomes very important for determining their final amounts. Mechanisms regulating translation can be targeted to specific cell cycle stages^18, 19, 20, 21^, as are pathways of RNA decay^22^. RNA decay and translation are controlled in part by the CNOT deadenylase complex, which has also been implicated in the regulation of the cell cycle^22, 23^. In GC B cells, post-transcriptional mechanisms mediated by microRNAs and RNA-binding proteins (RBPs) coordinate epigenetic remodelling, anabolic metabolism, survival and DNA repair mechanisms^23, 24, 25, 26, 27^. However, sequence-specific RBPs connected to the CNOT complex which coordinate the cell cycle and their role in GC B cells remain undefined.

ZFP36L1 and ZFP36L2 bind AU-rich elements within the 3’untranslated regions of mRNAs and are a subgroup of the many RBPs that promote RNA decay or repress translation via the CNOT1 complex^28^. They act as checkpoints for early development and effector function of B and T cells^29, 30, 31, 32, 33^, and show redundancy in multiple different cell contexts ^34, 35, 36, 37^. During B cell development, ZFP36L1 and ZFP36L2 are both required to maintain quiescence and facilitate VDJ rearrangement of the IgH chain^34^. Mice lacking *Zfp36l1* in B cells produced normal numbers of antigen-specific GC B cells following challenge yet display abnormal migration of antibody-secreting cells to the bone marrow^38^. If and how ZFP36L1 and ZFP36L2 have redundant or cooperative roles in B cells during the GC response is unknown. Here, we show that ZFP36L1 and ZFP36L2 are induced in response to antigen-sensing, promote survival and regulate cell cycle progression of GC B cells. By safeguarding the transition into mitosis, ZFP36L1 and ZFP36L2 protect GC B cells from replication stress.

## Results

### ZFP36L1 and ZFP36L2 follow independent expression kinetics in GC B cells

To generate expression-reporter mice we modified *Zfp36l1* and *Zfp36l2* by insertion of open reading frames encoding the fluorescent proteins mCherry or eGFP in frame with the start codon of ZFP36L1 and ZFP36L2, respectively (Supplementary Fig 1a, b). This design retains the transcriptional and post-transcriptional regulatory elements of the endogenous genes. In B cells, mCherry-ZFP36L1 and eGFP-ZFP36L2 fusion proteins are expressed at the expected molecular size, without breakdown products; for mCherry-ZFP36L1 the faster migrating band represents a phosphorylated form that is also evident in the endogenous protein (Supplementary Fig 1c, d). We traced the expression of mCherry-ZFP36L1 and eGFP-ZFP36L2 in B cells after intranasal infection with influenza A virus (HKx31). Both groups of reporter mice showed no difference in body weight loss or in the formation of GCs upon influenza infection (Supplementary Fig 1e, f). We analysed mCherry-ZFP36L1 and eGFP-ZFP36L2 expression in the mediastinal lymph node (medLN) and spleen at day 14 post-infection and found that both mCherry-ZFP36L1 and eGFP-ZFP36L2 were increased in GC B cells and decreased in plasmablasts, compared to naive B cells (Fig 1a, b and Supplementary Fig 1g, h). Although mCherry-ZFP36L1and eGFP-ZFP36L2 were both induced in GC B cells, the expression of eGFP-ZFP36L2 was similar between GC B cells with a LZ or DZ cell surface phenotype (Fig 1b and Supplementary Fig 1h). In contrast, mCherry-ZFP36L1 expression was induced more in LZ compared to DZ B cells (Fig 1a and Supplementary Fig 1g).

**Figure 1:**
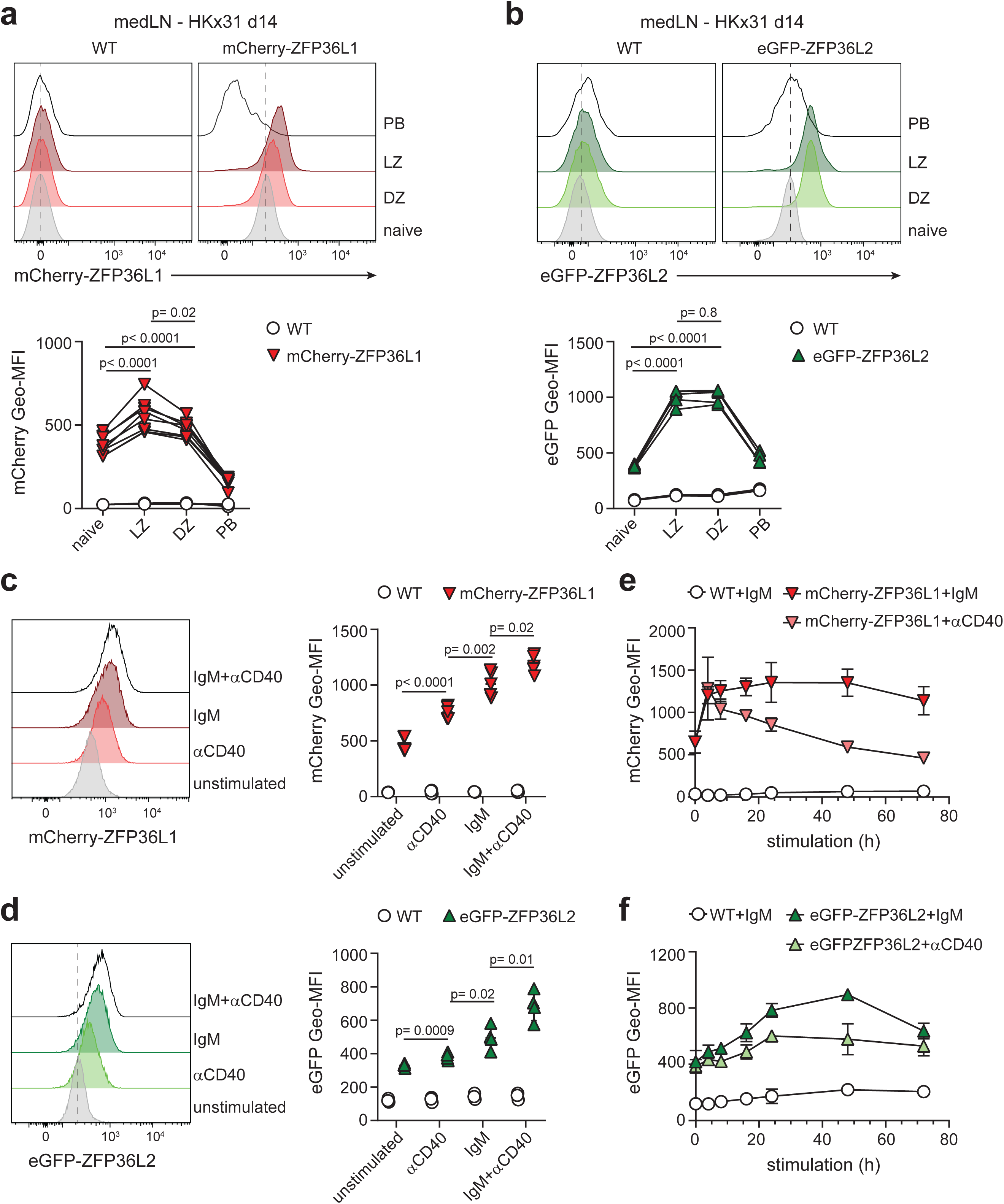
ZFP36L1 and ZFP36L2 follow independent expression kinetics in activated B cells. **a, b)** mCherry-ZFP36L1 **(a)** and eGFP-ZFP36L2 **(b)** reporter mice were infected i.n. with HKx31 influenza virus and medLNs were harvested 14 days later. mCherry or eGFP expression was evaluated in naive (CD19^+^ IgD^+^ CD38^+^ CD138^−^) cells, dark zone (DZ; CD19^+^ IgD^−^ CD38^−^ CD138^−^ CD95^+^ CXCR4^hi^ CD86^lo^) or light zone (LZ; CD19^+^ IgD^−^ CD38^−^ CD138^−^ CD95^+^ CXCR4^lo^ CD86^hi^) GC B cells, or plasmablasts (PB; CD19^int/lo^ IgD^−^ CD138^+^). Data were analysed using RM one-way ANOVA with Tukey’s multiple comparison test. **c, d)** mCherry-ZFP36L1 **(c)** and eGFP-ZFP36L2 **(d)** expression of B cells that were left unstimulated, or were stimulated for 24h with anti-CD40 antibody, anti-IgM F(ab)_2_ fragment, or a combination thereof. Unpaired Student’s *t*-test analysis was performed between indicated conditions. **e, f)** mCherry-ZFP36L1 **(e)** and eGFP-ZFP36L2 **(f)** expression kinetics at indicated time points, upon stimulation with anti-IgM or anti-CD40 antibody. **(a-f)** C57BL/6 (WT) mice were used as a negative control. Graphs display mean ±SD of 5-8 mice per group **(a, b)**, or 4-5 mice per group **(c, d)**, or 3-4 mice per group **(e, f)**. For each experimental setup, data were pooled from at least two independently performed experiments.

In the LZ, GC B cells are selected in an antigen- and T cell-dependent manner^39^. Because the expression of ZFP36L1 and ZFP36L2 is driven by B cell activation, we sought to elucidate the role of BCR triggering and/or T cell help in promoting their expression. We stimulated splenic B cells isolated from mCherry-ZFP36L1 and eGFP-ZFP36L2 reporter mice with soluble polyclonal F(abʹ)2 antibody specific for immunoglobulin M (anti-IgM), which we used to activate BCR signalling, or anti-CD40 antibody with IL-4, which we used to mimic T cell help. Both anti-IgM and anti-CD40 antibodies promoted and synergised with each other to induce expression of mCherry-ZFP36L1 and eGFP-ZFP36L2 (Fig 1c, d). Notably, the two RBPs followed different expression kinetics. mCherry-ZFP36L1 expression was rapid but transient, as it peaked within 4h of stimulation and declined over time (Fig 1e). In contrast, eGFP-ZFP36L2 levels steadily increased within the first 24h and reached a plateau independently of the type of stimulation (Fig 1f). Their different kinetics of expression upon B cell activation *in vitro* correlated with the different amounts of mCherry-ZFP36L1 and eGFP-ZFP36L2 we detected in *bona fide* GC B cells; whereas eGFP-ZFP36L2 was equally abundant in LZ and DZ B cells (Fig 1b), the expression of mCherry-ZFP36L1 was greatest in LZ B cells which have received recent BCR/CD40 stimulation (Fig 1a). Altogether, both ZFP36L1 and ZFP36L2 are produced by B cells in response to antigen-sensing and might act in different compartments of the GC according to their expression kinetics.

### ZFP36L1 and ZFP36L2 control the magnitude of GC responses

To study their function, we first used conditional knock-out mice in which a *loxP*-flanked region of the *Zfp36l1* or *Zfp36l2* gene was deleted by a transgene encoding Cre recombinase under the control of the CD2 locus control region^35^. In these mice, each RBP gene was deleted in early B cell development and B cell numbers were normal except for MZ B cells, which require ZFP36L1^40^. We immunized *Zfp36l1^fl/fl^* CD2^cre^ or *Zfp36l2^fl/fl^ CD2^cre^* mice with the thymus-dependent antigen 4-hydroxy-3-nitrophenylacetyl hapten conjugated to Keyhole Limpet Hemocyanin (NP-KLH) in alum and analysed the GC responses in the spleen 10 days later. Mice lacking ZFP36L1 or ZFP36L2 in lymphocytes had similar numbers of GC B cells (Fig 2a) and serum titres of NP-reactive IgG1 (Supplementary Fig 2a-b) compared to their littermate *Zfp36l1^fl/fl^* or *Zfp36l2^fl/fl^* control mice, indicating that individually these RBPs are not essential for GC responses.

**Figure 2:**
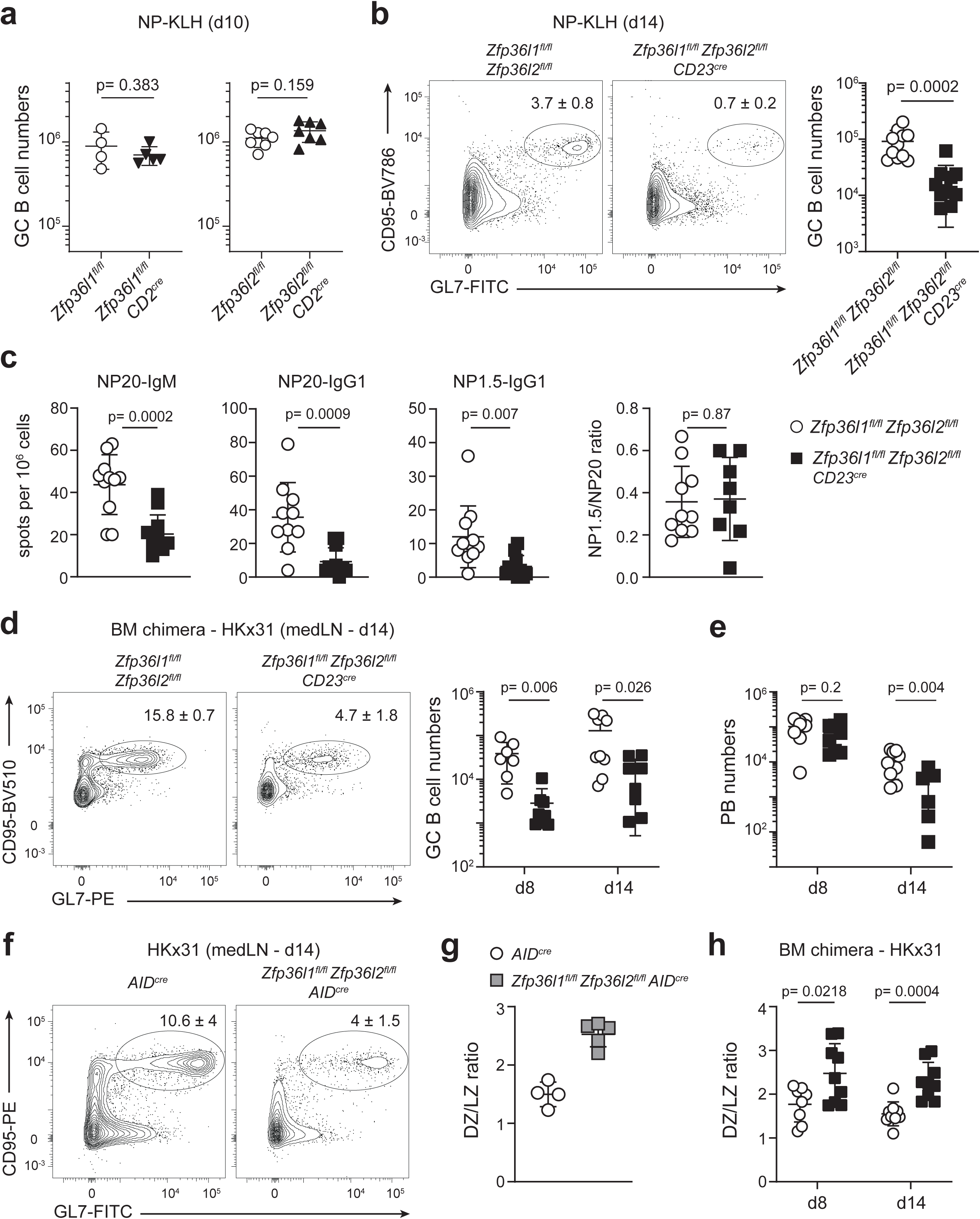
ZFP36L1 and ZFP36L2 control the magnitude of GC responses. **a)** GC B cell numbers in the spleen of *Zfp36l1^fl/fl^* CD2^cre^ mice or *Zfp36l2^fl/fl^ CD2^cre^* mice, and respective littermate controls, 10 days after immunization with NP-KLH. **b)** CD95 and GL7 expression identify GC B cells in *Zfp36l1^fl/fl^ Zfp36l2^fl/fl^ CD23^cre^* mice and littermate controls on day 14 after NP-KLH immunization. Cells were pre-gated as live CD19^+^ IgD^−^ cells. Numbers indicate mean frequency ±SD of 10-11 mice per group. Graph depicts corresponding absolute GC B cell numbers. **c)** ELISpot-analysis of splenic NP20-IgM, NP20-IgG1 and NP1.5-IgG1 antibody-secreting cells 14 days after NP-KLH immunization of *Zfp36l1^fl/fl^ Zfp36l2^fl/fl^* or *Zfp36l1^fl/fl^ Zfp36l2^fl/fl^ CD23^cre^* mice. NP1.5/NP20 IgG1 ratio was calculated. **d, e)** Chimeric mice were infected i.n. with HKx31 influenza virus and medLNs were analysed 8 or 14 days later for CD19^+^ IgD^−^ CD38^−^ CD95^+^ GL7^+^ GC B cell frequency and numbers **(d)**, and CD138^+^ TACI^+^ CD19^low/int^ IgD^−^ PB numbers **(e)**. **f)** Contour plots represent frequency of GC B cells in *Zfp36l1^fl/fl^ Zfp36l2^fl/fl^ AID^cre^* mice and *AID^cre^* controls upon HKx31 influenza infection. GC B cells were identified and displayed as in b. **g, h)** Ratio of CXCR4^hi^ CD86^low^ DZ versus CXCR4^low^ CD86^hi^ LZ GC B cells per mouse in *Zfp36l1^fl/fl^ Zfp36l2^fl/fl^ AID^cre^* versus *AID^cre^* control mice as in f **(g)**, or in chimeric mice as in d **(h)**. **c, d, f)** Unpaired Student’s *t*-test analysis; n= 11 mice per group **(b, c)**; n= 7-9 mice per group **(d, e, h)**; n= 4-5 mice per group **(f, g)**. In each graph, data were pooled from at least two independently performed experiments.

ZFP36L1 and ZFP36L2 have redundant roles during B cell development, and their early deletion using CD2^cre^ causes T lymphoblastic leukaemia in mice^34, 35^ making this system unsuitable to study B cell responses. To elucidate the intrinsic role of ZFP36L1 and ZFP36L2 in mature B cells, we crossed *Zfp36l1^fl/fl^ Zfp36l2^fl/fl^* mice with transgenic mice expressing Cre recombinase under the direction of the low-affinity IgE receptor FcεRII/CD23 promoter (CD23^cre^), which causes deletion at the transitional 2 (T2) B cell stage^41^. We immunized *Zfp36l1^fl/fl^ Zfp36l2^fl/fl^* (control) or *Zfp36l1^fl/fl^ Zfp36l2^fl/fl^ CD23^cre^* littermate mice with NP-KLH in alum and found that loss of both *Zfp36l1* and *Zfp36l2* led to five-fold reduction in the proportion and absolute numbers of GC B cells (Fig 2b). Enumeration of splenic NP-specific antibody secreting cells by ELISpot revealed a reduction in both IgM- and IgG1-secreting cells, irrespective of antibody affinity (Fig 2c). The ratio between NP-1.5-binding (high affinity) and NP-20-binding (high + low affinity) IgG1-secreting cells was comparable between *Zfp36l1^fl/fl^ Zfp36l2^fl/fl^ CD23^cre^* and control mice (Fig 2c). This indicates that, although the affinity maturation of GC B cells can take place in the absence of ZFP36L1 and ZFP36L2, the overall magnitude of the GC response is impaired.

To assess whether the reduction in GC B cell numbers was due to a cell-autonomous defect in B cell activation, we generated mixed bone marrow (BM) chimeras by reconstituting irradiated B6.SJL mice with a 1:1 mixture of BM cells from CD45.1^+^ B6.SJL mice and BM cells from CD45.2^+^ *Zfp36l1^fl/fl^ Zfp36l2^fl/fl^ CD23^cre^* or CD45.2^+^ *Zfp36l1^fl/fl^ Zfp36l2^fl/fl^* mice as a control group. Chimeric mice were intranasally infected with HKx31 influenza virus, and medLNs and spleens were analysed at day 8 and 14 post-infection, to capture the formation and expansion of GCs, respectively. In the absence of ZFP36L1 and ZFP36L2, the proportion and numbers of GC B cells were significantly reduced at both time points (Fig 2d and Supplementary Fig 2c). Whereas plasmablast (PB) formation was also reduced at day 14 post-infection, PB numbers measured at day 8 were comparable between the two genotypes (Fig 2e), suggesting an intact extrafollicular response. Thus, *Zfp36l1-Zfp36l2-*deletion drives an intrinsic defect in GC B cells, which might occur after their initial activation.

To address whether the reduction in GC size was due to the inability to expand and/or maintain GC B cells, rather than impaired B cell activation, we generated *Zfp36l1^fl/fl^ Zfp36l2^fl/fl^ AID^cre^* mice, in which the Cre recombinase is expressed under the control of an activation-induced cytidine deaminase (*Aicda*) promoter^41^, thus resulting in *Zfp36l1* and *Zfp36l2* deletion after antigen activation of B cells. As observed with CD23^cre^ mediated deletion, the percentage of GC B cells were reduced by about three-fold in *Zfp36l1^fl/fl^ Zfp36l2^fl/fl^ AID^cre^* mice compared to *AID^cre^* control mice 14 days after influenza infection (Fig 2f). The magnitude of GC responses was also impaired when *Zfp36l1^fl/fl^ Zfp36l2^fl/fl^ AID^cre^* mice were immunized with NP-KLH compared to *Zfp36l1^fl/fl^ Zfp36l2^fl/fl^* littermates, which we used as control mice (Supplementary Fig 2d). We further characterised GC B cells based on cell surface expression of markers for DZ and LZ cells and found an increase in DZ/LZ ratio for *Zfp36l1*-*Zfp36l2*-deficient GC B cells in both the AID^cre^ and CD23^cre^ systems (Fig 2g, h and Supplementary Fig 2e, f). Altogether, our data show that in B cells ZFP36L1 and ZFP36L2 are not essential for affinity maturation but are required to promote the cellularity of the GC response.

### Deletion of ZFP36L1 and ZFP36L2 causes accumulation of GC B cells in G2-M phase and increases cell death

To investigate whether ZFP36L1 and ZFP36L2 were required for the expansion of GC B cells, we analysed their cell division cycle *in vivo* by measuring incorporation of the thymidine analogue bromodeoxyuridine (BrdU) and DNA content (7AAD). Upon influenza infection, the proportion of GC B cells in G0-G1 phase (BrdU^−^ 7AAD^−^) or in S phase (BrdU^+^) was similar between *Zfp36l1^fl/fl^ Zfp36l2^fl/fl^ AID^cre^* mice and *AID^cre^* control mice. However, *Zfp36l1*-*Zfp36l2*-deficient GC B cells had a greater proportion of cells in G2-M phase (BrdU^−^ 7AAD^+^) than in control mice (Fig 3a). Accumulation of *Zfp36l1*-*Zfp36l2*-deficient GC B cells in G2-M phase was found in cells with both a DZ or a LZ phenotype (Fig 3b) and confirmed in competitive chimeras (Supplementary Fig 3a), thus showing that this was a cell-autonomous effect. Because the difference in cell cycle progression did not directly explain the reduction of GC B cell numbers, we next assessed whether ZFP36L1 and ZFP36L2 were also required to control cell death. We measured the expression of active-caspase 3 as a marker of early apoptosis and found that the proportion of active-Casp3^+^ GC B cells was increased four-fold upon deletion of ZFP36L1 and ZFP36L2 (Fig 3c and Supplementary Fig 3b). Although observed in both compartments, the increase in cell death was more pronounced in LZ B cells (4-fold increase), than in DZ B cells (2-fold increase) (Fig 3d). Together, these data show that ZFP36L1 and ZFP36L2 control progression into mitosis and protect GC B cells from apoptosis.

**Figure 3:**
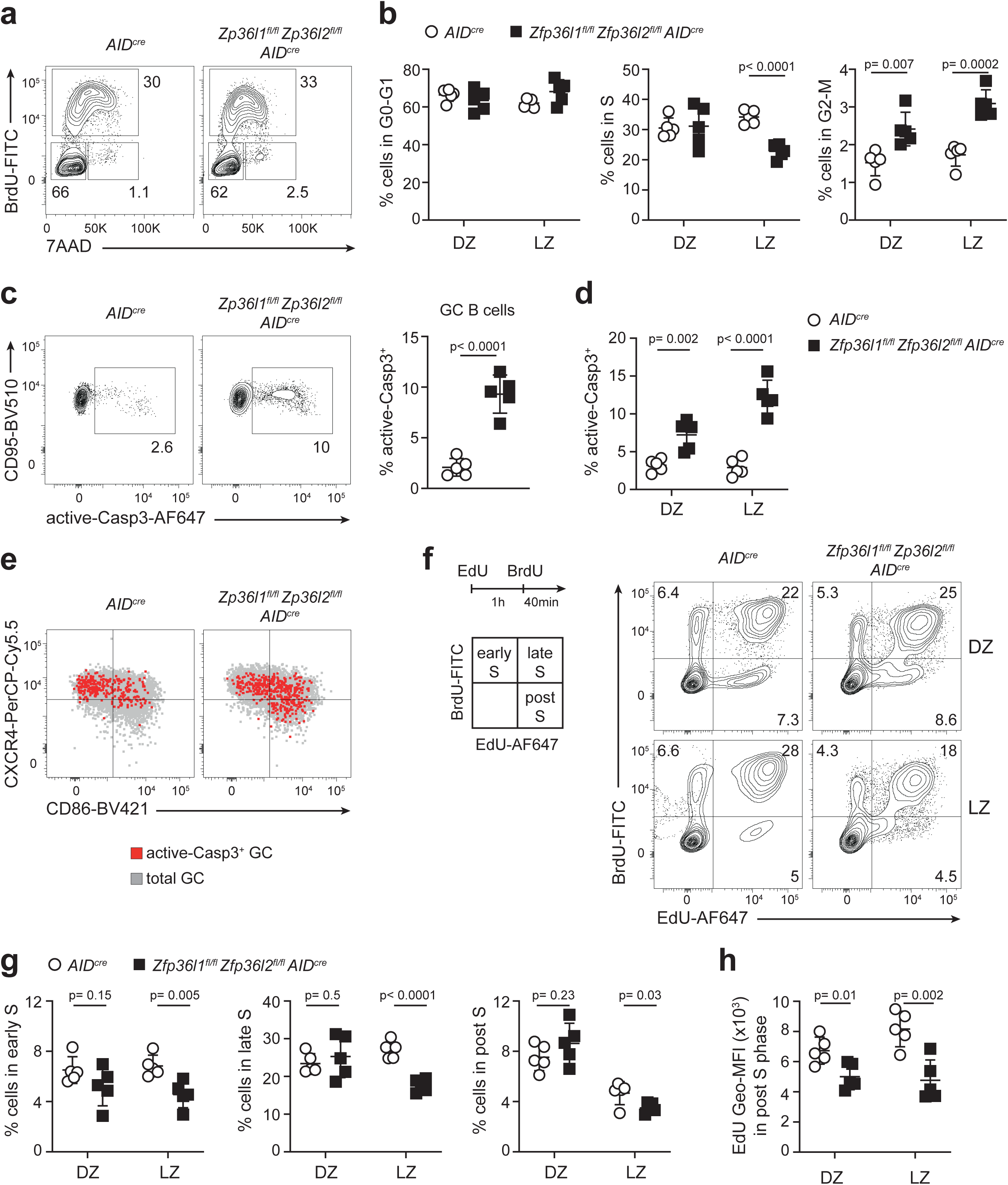
ZFP36L1 and ZFP36L2 control cell cycle progression and apoptosis of GC B cells. *Zfp36l1^fl/fl^ Zfp36l2^fl/fl^ AID^cre^* mice or *AID^cre^* control mice were analysed 14 days after i.n. HKx31 influenza virus infection. **a)** GC B cells (CD19^+^ CD38^−^ CD95^+^ GL7^+^) isolated from medLNs were analysed at various stages of cell cycle based on flow cytometry staining for BrdU and the membrane-impermeable DNA-intercalating dye 7-AAD. Numbers adjacent to outlined areas indicate frequency of cells in each phase. **b)** Graphs display frequencies of DZ or LZ B cells in G0-G1, S, or G2-M phase of the cell cycle as identified in panel a. **c)** Representative contour plot and compilation of apoptotic GC B cells, evaluated based on active-Caspase 3 staining. **d)** Percentage of active-Casp3^+^ cells within CXCR4^hi^ CD86^low^ DZ and CXCR4^low^ CD86^hi^ LZ B cells. **e)** Overlay dot plots depict DZ and LZ distribution of total GC B cells (grey) or active-Casp3^+^ GC B cells (red). **f-g)** EdU and BrdU sequential labelling to identify DZ and LZ GC B cells in early (EdU^−^ BrdU^+^), late (EdU^+^ BrdU^+^) or post (EdU^+^ BrdU^−^) S phase of the cell cycle. Representative contour plots in (**f**), compilation in (**g**). **h)** Graph depicts EdU geometric median fluorescence intensity (Geo-MFI) in DZ or LZ GC B cells within post S phase. **(a-h)** Graphs display mean ±SD of 5 mice per group. Data represent at least two independently performed experiments and were analysed with an unpaired Student’s *t*-test.

During positive selection, a fraction of GC B cells evades apoptosis and undergoes a first cell division cycle within the LZ. Subsequently, this selected LZ B cell subpopulation migrates to the DZ, where it clonally expands^7, 13^. We found that, in addition to the accumulation of LZ and DZ B cells in the G2-M phase, the proportion of LZ B cells in S phase was reduced in *Zfp36l1^fl/fl^ Zfp36l2^fl/fl^ AID^cre^* mice compared to *AID^cre^* control mice (Fig 3b). This observation could be explained by the concomitant increase of active-caspase-3 positive *Zfp36l1*-*Zfp36l2*-deficient LZ B cells (Fig 3d, e), indicating that the proportion of LZ B cells re-entering cell cycle upon positive selection was also controlled by ZFP36L1 and ZFP36L2.

Next, we examined whether accumulation in G2-M phase was due to different speed of progression through S phase. For this purpose, we pulsed influenza-infected mice first with the thymidine analogue 5-ethynyl-2′-deoxyuridine (EdU), and 1h later with BrdU. At 40 minutes after BrdU administration, we identified early S (EdU^−^ BrdU^+^), late S (EdU^+^ BrdU^+^), and post S phase (EdU^+^ BrdU^−^) within LZ and DZ B cells (Fig 3f, g). In agreement with our finding using BrdU alone (Fig 3b), the proportion of DZ B cells in early, late and post S phase was comparable between *Zfp36l1^fl/fl^ Zfp36l2^fl/fl^ AID^cre^* mice and *AID^cre^* control mice (Fig 3f, g). Instead, the entry and progression of *Zfp36l1*-*Zfp36l2*-deficient LZ B cells through the S phase was reduced by 1.5-fold compared to control (Fig 3f, g), confirming an impairment in their ability to re-enter cell cycle. When analysing the EdU/BrdU incorporation profile (Fig 3f), we observed that *Zfp36l1*-*Zfp36l2*-deficient cells in post-S-phase showed reduced EdU intensity compared to control (Fig 3h). Considering that cells in post-S-phase have completed DNA replication and are moving into G2, *Zfp36l1*-*Zfp36l2*-deficient cells displaying lower EdU intensity appear to have entered G2-M faster than controls. Altogether, our data show a dual role for ZFP36L1 and ZFP36L2: 1) in the LZ they contribute to positive selection by promoting initiation of the S phase; 2) upon DNA replication, they regulate the progression of both DZ and LZ B cells into mitosis and protect them from apoptosis.

### ZFP36L1 and ZFP36L2 control the G2-M checkpoint of GC B cells

To elucidate the molecular mechanisms by which ZFP36L1 and ZFP36L2 control cell cycle progression, we used an *in vitro* GC B cell culture system^42^ that recapitulated the accumulation of cells in the G2-M phase observed in *bona fide* GC B cells (Fig 4a) and enabled the generation of sufficient numbers of GC-like B cells for molecular studies. To identify RNA targets of ZFP36L1, we performed individual-nucleotide-resolution crosslinking immunoprecipitation (iCLIP) using cytoplasmic extracts of GC-like B cells derived from mCherry-ZFP36L1 reporter mice. Immunoprecipitation of the mCherry-ZFP36L1 fusion protein confirmed that ZFP36L1 preferentially binds to AU-rich elements within the 3’untranslated region (3’UTR) of mature transcripts (Supplementary Fig 4a). ZFP36L1-bound transcripts in GC-like B cells were enriched for cell cycle-related genes, which were further classified based on the presence of AU-rich elements within their 3’UTR (Fig 4b and Supplementary Table 1). By pairing iCLIP with global transcriptomic analysis, we found that the abundance of eight cell cycle-related RNA targets (*Ccnd2, Ccne2, Cdkn1a, Cep76, Csnk1e, Mdm2, Nup88, Ska2*) was increased in *Zfp36l1*-*Zfp36l2*-deficient GC-like B cells compared to control (Fig 4b and Supplementary Table 1), consistent with the direct binding of these RBPs promoting mRNA decay. However, the abundance of 33 cell cycle-related RNA targets was comparable between *Zfp36l1^fl/fl^ Zfp36l2^fl/fl^ CD23^cre^* and *Zfp36l1^fl/fl^ Zfp36l2^fl/fl^* GC-like B cells, consistent with ZFP36L1 mediating functions in addition to promoting RNA decay such as the regulation of mRNA translation^43, 44^ (Fig 4b and Supplementary Table 1).

**Figure 4:**
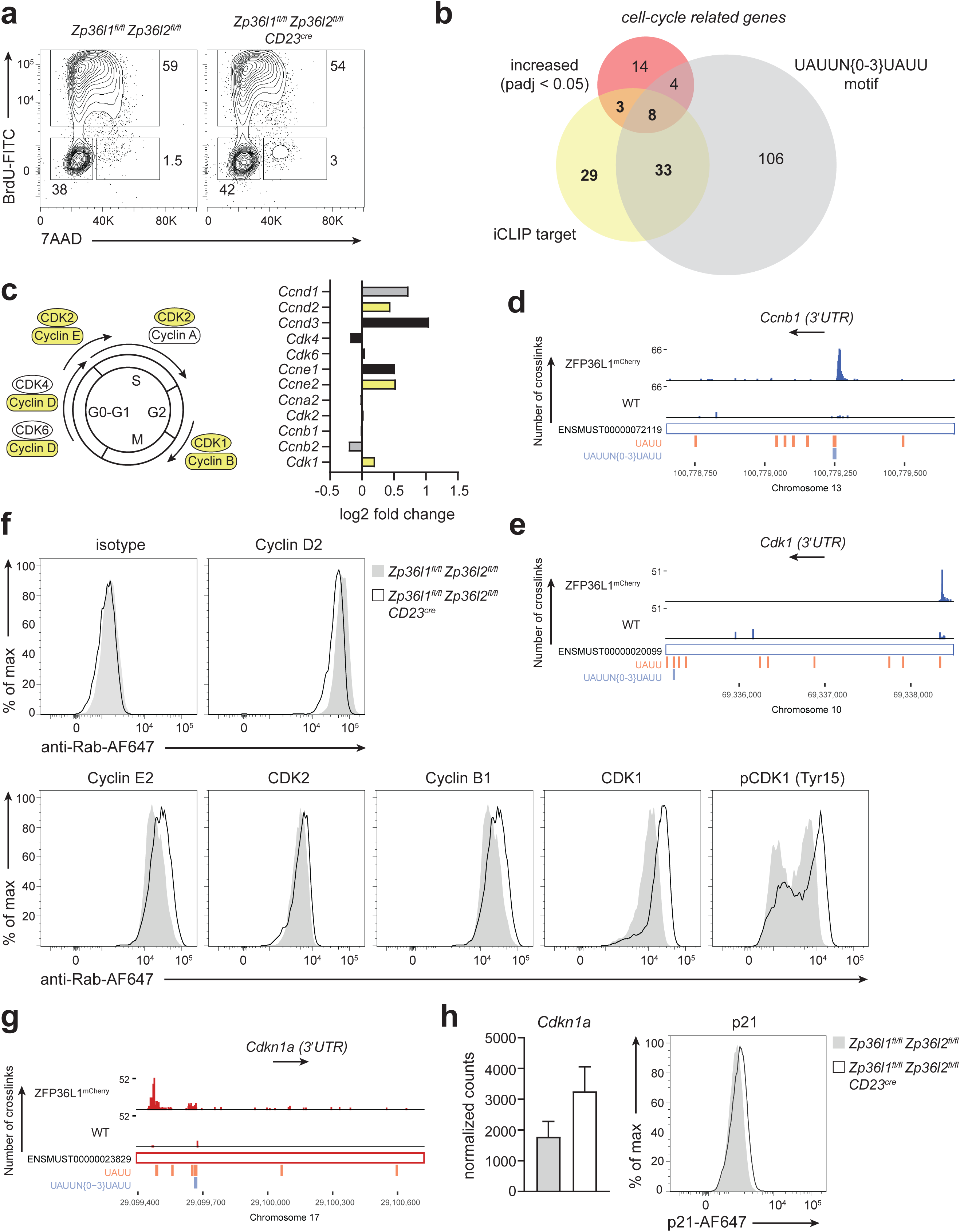
ZFP36L1 and ZFP36L2 control the G2-M checkpoint of GC-like B cells. B cells isolated from LNs of *Zfp36l1^fl/fl^ Zfp36l2^fl/fl^ CD23^cre^* mice or *Zfp36l1^fl/fl^ Zfp36l2^fl/fl^* control mice were co-cultured for 7 days in the presence of CD40LB-expressing feeder cells. **a)** BrdU and 7AAD staining of CD95^+^ CD138^−^ GC-like B cells. **b)** Venn diagram for genes within the Reactome “Cell cycle” gene set, showing the overlap between those that are increased in *Zfp36l1^fl/fl^ Zfp36l2^fl/fl^ CD23^cre^* GC-like B cells compared to *Zfp36l1^fl/fl^ Zfp36l2^fl/fl^* control cells, bound by mCherry-ZFP36L1 in their 3’UTR, or contain two UAUU motifs separated by up to three nucleotides within their 3’UTR. **c)** Left: schematic representation of cyclins and CDKs regulating cell cycle progression; iCLIP targets in yellow. Right: Log2 fold change (KO/control) of mRNA encoding cyclins and CDKs as measured by RNA-seq. In yellow iCLIP targets encompassing UAUUN{0-3}UAUU motif within their 3’UTR; in grey mRNAs containing AU-rich binding motifs but not identified as iCLIP targets; in black mRNA without UAUUN{0-3}UAUU binding motif. **d, e, g)** iCLIP data from GC-like B cells (d7) showing reads across *Ccnb1* **(d)**, *Cdk1* **(e)**, *Cdkn1a* **(g)** transcripts with an expanded view of the 3’UTR. **f)** Histograms depict protein expression of indicated cell-cycle related genes. n= 3 mice per group; data representative of at least two independently performed experiments. **h)** Left: graph displays normalized counts (mean±SD) of *Cdkn1a* as measured by RNAseq. Right: protein expression as measured by flow cytometry. n= 3 mice per group.

Among the candidate ZFP36L1-targets for which mRNA abundance was unchanged in *Zfp36l1*-*Zfp36l2*-deficient GC-like B cells, were the mRNAs encoding cyclin B1 and CDK1, which together are essential to enable the transition from G2 into mitosis (Fig 4c). *Ccnb1* and *Cdk1* mRNAs were enriched with high confidence by iCLIP, based on the number of measured crosslinking sites (Fig 4d, e) but their mRNA amounts were not different (Fig 4c - right panel). We therefore used flow cytometry to assess whether ZFP36L1-binding had functional consequences on protein abundance. In parallel to cyclin B1 and CDK1, we measured protein levels of cyclin D2, cyclin E2 and CDK2, which were also identified by iCLIP as ZFP36L1-bound mRNAs (Supplementary Fig 4b-d). Consistent with increased mRNA levels (Fig 4c - right panel), *Zfp36l1^fl/fl^ Zfp36l2^fl/fl^ CD23^cre^* GC-like B cells expressed more cyclin E2 protein compared *Zfp36l1^fl/fl^ Zfp36l2^fl/fl^* control cells, however this was not the case for cyclin D2 (Fig 4f and Supplementary Fig 4e). While *Cdk2* mRNA and protein amounts were unchanged (Fig 4c - right panel, f), *Zfp36l1*-*Zfp36l2*-deficient GC-like B cells expressed more cyclin B1 and CDK1 protein than *Zfp36l1^fl/fl^ Zfp36l2^fl/fl^* control cells (Fig 4f and Supplementary Fig 4e), thus indicating a role for ZFP36L1 and ZFP36L2 in regulating progression through G2-M phase of GC B cells.

Cyclin B1 and CDK1 form complexes during the S and G2 phases, and their activity is regulated through phosphorylation of CDK1^45, 46^. We found that *Zfp36l1*-*Zfp36l2*-deficient GC-like B cells displayed increased phosphorylation on the inhibitory Tyr-15 residue of CDK1 (Fig 4f and Supplementary Fig 4e), which promotes accumulation of inactive cyclin B1-CDK1 complexes and cell cycle arrest in G2^46^. Retention of inactive cyclin B1-CDK1 complexes is at least in part mediated by p21^47^ (encoded by *Cdkn1a*), which is also a target of ZFP36L1 (Fig 4g) and more abundant in *Zfp36l1*-*Zfp36l2*-deficient GC-like B cells both at the mRNA and protein level (Fig 4h and Supplementary Fig 4f). Altogether, our data show that ZFP36L1 and ZFP36L2 control the completion of mitosis by repressing the expression and function of a cell cycle-related RNA post-transcriptional regulon, which includes both positive and negative regulators of the cell cycle. By doing this, ZPF36L1 and ZFP36L2 become essential components of the G2-M checkpoint of GC B cells.

### ZFP36L1 and ZFP36L2 protect GC B cells from replication stress

The G2-M checkpoint ensures that cells with damaged DNA are eliminated and do not complete mitosis. Replication stress during S phase is a major cause of DNA damage and is generated by slowing or stalling of the replication fork during DNA synthesis^48^. Upon selection, GC B cells accelerate their proliferation rate by controlling replication fork progression^4^. We investigated whether replication fork progression was hampered in GC B cells lacking ZFP36L1 and ZFP36L2, thus causing replication stress. For this purpose, we performed Transferase-Activated End Ligation sequencing (TrAEL-seq^49^) on GC-like B cells derived from *Zfp36l1^fl/fl^ Zfp36l2^fl/fl^ CD23^cre^* mice and *Zfp36l1^fl/fl^ Zfp36l2^fl/fl^* control mice. TrAEL-seq labels the free 3’ end of the leading strand at DNA replication forks, thereby determining both replication fork abundance and direction of travel in unsynchronised cell populations^49^. Defects in DNA replication fork progression result in the accumulation of replication forks around replication initiation zones, which is detectable through an increase in TrAEL-seq read count^50^. In *Zfp36l1*-*Zfp36l2*-deficient GC-like B cells the directionality of replication fork progression was unchanged (Supplementary Fig 5a), however TrAEL-seq read counts around active replication initiation zones were increased compared to control (Fig 5a), indicating that DNA replication forks aberrantly accumulate around active replication initiation zones in *Zfp36l1*-*Zfp36l2*-deficient GC-like B cells. Notably, even control cells show an accumulation of replication forks around replication initiation zones, indicating that the extremely rapid proliferation of GC B cells exerts replication stress that is accentuated in *Zfp36l1*-*Zfp36l2*-deficient GC-like B cells (Fig. 5a). Eukaryotic cells fire replication origins in a regulated fashion, forming early- and late-replication initiation zones^51^. Interestingly, we found that in *Zfp36l1*-*Zfp36l2*-deficient GC-like B cells the replication fork distribution was skewed away from late-replicating regions (Fig 5b), but there was no evidence that stalling around early replicating initiation zones was compensated by the firing of additional origins in late-replicating regions (Fig 5b, c). These results suggest that the stalling occurs in early S phase and remains unresolved in G2-M phase. Because failure to resolve stalled replication forks results in an excess of single-stranded DNA, we measured the phosphorylated form of the histone variant H2AX (pH2AX), which marks single-stranded DNA and is phosphorylated in response to replication stress^52^. Control GC-like B cells (*Zfp36l1^fl/fl^ Zfp36l2^fl/fl^*) displayed high pH2AX levels during DNA replication in S phase, and low pH2AX levels upon transition to G2-M phase (Fig 5c). In contrast, *Zfp36l1^fl/fl^ Zfp36l2^fl/fl^ CD23^cre^* GC-like B cells maintained high pH2AX levels upon transition to G2-M phase (Fig 5d), showing that *Zfp36l1*-*Zfp36l2*-deficient GC-like B cells that were arrested in G2-M phase were vulnerable to DNA breaks.

**Figure 5:**
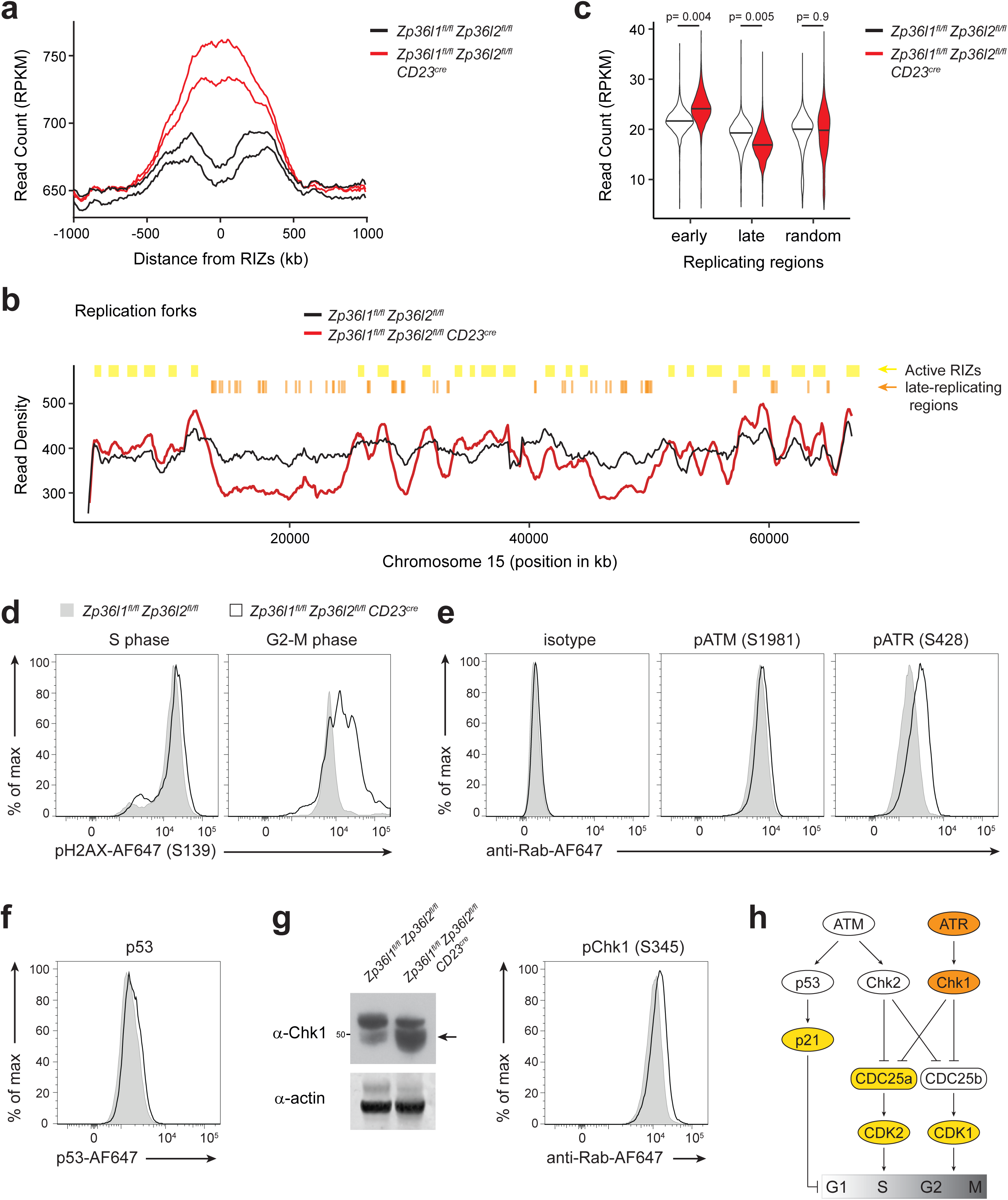
ZFP36L1 and ZFP36L2 protect GC-like B cells from replication stress. **a)** Quantitation trend plot of read counts for TrAEL-seq data centred around active replication initiation zones (RIZs), for *Zfp36l1^fl/fl^ Zfp36l ^fl/fl^* control (black) and Zfp36l *^fl/fl^* Zfp36l2*^fl/fl^ CD23^cre^* (red) GC-like B cells (d7). Graph displays two independently performed experiments (n= 2 mice per group). **b)** Representative read density plot for of *Zfp36l1^fl/fl^ Zfp36l2^fl/fl^* control (black) and *Zfp36l1^fl/fl^ Zfp36l2^fl/fl^ CD23^cre^* (red) over a randomly selected region of the genome (chromosome 15). Active replication initiation zones (RIZs) are indicated in yellow; late-replicating regions are indicated in orange Shown TrAEL-Seq data are an average of two biological replicates. c) Graphs depicts TrAEL-seq read counts that were detected around early replication initiation zones, late replication initiation zones, or randomly across the genome. **d)** Representative histogram of pH2AX expression in *Zfp36l1^fl/fl^ Zfp36l2^fl/fl^ CD23^cre^*(black line) and *Zfp36l1^fl/fl^ Zfp36l2^fl/fl^* control (grey) GC-like B cells (d7) that were in S phase (BrdU^+^) or in G2-M phase (BrdU^−^ 7AAD^+^). **e, f, g)** Histograms depict protein expression or phosphorylation of factors mediating DNA damage responses, as depicted by flow cytometry. **f)** Western blot of CHK1 and actin. n= 3 mice per group; data representative of at least two independently performed experiments. **g)** Schematic representation of DNA damage responses that control cell cycle progression. In yellow ZFP36L1 iCLIP targets; in red factors that do not directly interact with ZFP36L1, but show increased activity in *Zfp36l1^fl/fl^ Zfp36l2^fl/fl^ CD23^cre^* GC-like B cells.

Signals elicited by aberrant DNA replication activate DNA damage responses, which are primarily mediated by ATM and ATR kinases. ATR is central to the replication stress response and, upon activation, promotes cell cycle arrest while suppressing the firing of late-replication initiation zones^53^. In agreement with this, *Zfp36l1^fl/fl^ Zfp36l2^fl/fl^ CD23^cre^* GC-like B cells displayed increased phosphorylation of ATR, but not ATM kinase (Fig 5e). In addition, whereas the expression of ATM-dependent p53 was unchanged (Fig 5f), total CHK1 protein and its phosphorylation on the ATR-dependent residue Ser345 were increased in *Zfp36l1*-*Zfp36l2*-deficient GC-like B cells compared to control (Fig 5g). In conclusion, we show that GC B cells undergoing replication stress accumulates in G2-M phase when ZFP36L1 and ZFP36L2-mediated post-transcriptional regulation is lost. These cells display increased activity of the ATR/CHK1 DNA damage response, which in turn controls cell cycle progression and survival of GC B cells (Fig 5h).

## Discussion

In this study, we show that the RBPs ZFP36L1 and ZFP36L2 coordinate the expression and function of a cell cycle-related RNA post-transcriptional regulon that is essential to control expansion and maintenance of GC B cells. In the LZ, GC B cells express ZFP36L1 and ZFP36L2 in response to antigen- and CD40-mediated signals to protect them from apoptosis and promote cell cycle re-entry. In both LZ and DZ B cells, ZFP36L1 and ZFP36L2 safeguard the successful completion of mitosis by regulating the balance between CDK1 and p21-mediated negative feedback of the cell cycle progression. CDK1, in addition to controlling the cell cycle, is required for faithful completion of DNA replication, DNA damage repair and checkpoint activation^54, 55, 56, 57^. In particular, CDK1 can promote excessive firing of replication origins, which depletes nucleotides and triggers replication fork stalling^54, 58^. We found that in the absence of ZFP36L1 and ZFP36L2, GC B cells that accumulate in G2-M phase exhibit replication stress with stalling of DNA replication forks at early replicating regions, which are not resolved in late S phase. Despite this, *Zfp36l1*-*Zfp36l2*-deficient GC B cells progressed faster to G2-M phase, where many were arrested and eventually died by apoptosis. It is therefore possible that ZFP36L1 and ZFP36L2, by limiting CDK1 activity, could promote replication fork progression in GC B cells while preserving the nucleotide pool for proper DNA replication. This hypothesis is however difficult to elucidate since current technologies for mechanistic studies of DNA replication are restricted to *in vitro* culture systems, which still imperfectly recapitulate the complexity of the GC environment.

Studies of the cell division cycle and replication stress in other cell types suggest that the balance of CDK1 activity and p21 in mother cells influences the cell division cycle of the daughter cell^21, 59, 60^. We show evidence that p21 is elevated in GC B cells lacking ZFP36L1 and ZFP36L2, but it is insufficient to block mitosis, thus daughter cells may be born with greater amounts of DNA damage arising from replication stress. In this scenario, GC B cells in transit between DZ and LZ might not have had sufficient time to adequately repair damaged DNA before selection and cell cycle re-entry in the LZ. Whether or not the amounts of replication stress sustained in the previous round of cell division influences the establishment of quiescence upon LZ re-entry and thus the capacity for initiating new cell cycles remains unexplored. ZFP36L1 and ZFP36L2 thus emerge to be important for GC B cell survival by coupling antigen receptor- and CD40-mediated signals to rapid cell cycling and somatic mutation.

In developing B and T cells ZFP36L1 and ZFP36L2 control the cell cycle by enforcing quiescence^34, 36^, but unlike GC B cells there is no evidence for a role in the G2-M checkpoint. These differences may reflect distinct regulation of the cell cycle in GC B cells and in response to pre-TCR and pre-BCR signals. In both cell contexts, the cell cycle is integrated with the DNA damage response, but the extent of DNA mutation in GC B cells undergoing SHM is greater than that of developing lymphocytes undergoing VDJ recombination. Furthermore, the speed of cell division is faster in GC B cells than in developing B and T cells^61, 62^. This increases the vulnerability of GC B cells to accumulating transforming mutations.

Tandem C3H-zinc-finger containing RBPs related to ZFP36L1 and ZFP36L2 were found to be regulators of the cell cycle in yeasts^63, 64^, suggesting that the ZFP36 family have coevolved with the cell cycle over an extended period. ZFP36L1 and ZFP36L2 are in fact deeply embedded in the regulation of the cell division cycle. This is evident from our finding that ZFP36L1 interacts with mRNAs that both promote and restrain the cell cycle^34, 65, 66^. While the ability to inhibit both positive and negative regulators of the cell cycle might appear paradoxical, they are consistent with the properties of an integrator (in this case the RBPs) that can coordinate the timing and location of expression of multiple regulatory proteins^67^. For example, the ability of p21 to negatively regulate the later stages of the cell cycle must be balanced against a role at earlier stages where it can promote stability of Cyclin/CDK complexes^68^, and it is possible that ZFP36L1 and ZFP36L2 are part of this balancing mechanism. Furthermore, as protein translation rates integrate past mitogenic signals of mother cells to control the proliferation-quiescence decision in daughter cells^21^, ZFP36L1 and ZFP36L2 may operate at distinct cell cycle stages to control translation of proteins.

Consistent with this, ZFP36L1 and ZFP36L2 expression and subcellular localization have been shown to be regulated in a cell cycle-dependent manner. Whereas ZFP36L1 accumulates in the nuclei of cells arrested in G1-S phase^69^, both ZFP36L1 and ZFP36L2 are localized in the cytoplasm when cells undergo mitosis^70, 71^. It is possible that, depending on their subcellular location, ZFP36L1 and ZFP36L2 target different mRNAs and thus control specific phases of the cell cycle. If this was the case, the apparent redundancy between these paralogs could result from a synthetic interaction. This hypothesis however warrants further investigation as a limitation of the study is the inability to identify direct RNA-targets for ZFP36L2 due to lack of specificity and sensitivity of affinity reagents. In addition to this, the activity of these RBPs is strictly regulated through phosphorylation. For example, ZFP36L1 is phosphorylated upon activation of the p38/MK2 pathway which is triggered downstream of ATM and ATR in response to DNA damage. MK2 phosphorylates ZFP36L1 *in vitro*^72^ to reduce its association with the RNA decay machinery and thus promote stabilisation of some of its targets. MK2 is also required to prolong G2-M checkpoint maintenance^73^; whether this occurs through modulation of ZFP36L1 and ZFP36L2 activity in GC B cells remains however unknown.

In conclusion, we show that ZFP36L1 and ZFP36L2 control cell cycle progression and the G2-M checkpoint of GC B cells by regulating the activity of a cell-cycle related RNA post-transcriptional regulon. This provides a safeguarding mechanism to GC B cells that is required to protect them from replication stress and maintain genome integrity. How ZFP36L1 and ZFP36L2 contributes to genome instability and malignant transformation remain to be examined.

## Methods

### Mice

Floxed alleles of *Zfp36l1* or *Zfp36l2* allele^35^ were crossed with Tg(*CD2-cre*)4^Kio74^ , Tg(*Fcer2a-cre*)5^Mbu^ (*CD23^cre^*) or Tg(*Aicda-cre*)9^Mbu^ (*AID^cre^*) ^41^ to generate the mice used in this study. mCherry-ZFP36L1 and eGFP-ZFP36L2 knock-in reporter mice were generated by Cyagen bioscience and designed to harbour the sequence encoding mCherry or eGFP without stop codon, in-frame upstream of the ATG translation start site of *Zfp36l1* or *Zfp36l2*, respectively. The *Zfp36l1* targeting vector featured diphtheria toxin A, a neomycin resistance cassette flanked by FRT recombination sequences and homology arms, which were amplified from a mouse genomic BAC clone. The *Zfp36l2* targeting vector contained self-deletion anchor sites. When homozygous, we did not observe pathology in the reporter mice indicative of loss of function^43, 75^.

All mouse strains listed above were maintained on a C57BL/6 (B6-CD45.2) background. B6-CD45.2 mice and B6.SJL-*Ptprc^a^Pepc^b^*/BoyJ (B6.SJL-CD45.1) mice were bred and housed at the Biological Support Unit (BSU) of the Babraham Institute (UK). Since the opening of the Babraham BSU in 2009, no primary pathogens or additional agents listed in the FELASA recommendations have been confirmed during health monitoring surveys of the stock holding rooms. Ambient temperature was ∼19-21°C and relative humidity 52%. Lighting was provided on a 12-hour light: 12-hour dark cycle including 15 min ‘dawn’ and ‘dusk’ periods of subdued lighting. After weaning, mice were transferred to individually ventilated cages with 1-5 mice per cage. Mice were fed CRM (P) VP diet (Special Diet Services) ad libitum and received seeds (e.g. sunflower, millet) at the time of cage-cleaning as part of their environmental enrichment. All mouse experiments were approved by the Babraham Institute Animal Welfare and Ethical Review Body and complied with existing European Union and United Kingdom Home Office legislation and local standards.

8 to 12-week-old female or male mice were used for both *in vivo* and *in vitro* experiments. Although no sex-associated differences were observed, control mice were sex- and aged-matched littermates negative for a flox-carrying allele or a Cre-carrying allele. Sex- and aged-matched C57BL/6 (WT) mice were used as a control for MCherry-ZFP36L1and EGFP-ZFP36L2 reporter mice.

### Animal procedures

For generation of competitive bone marrow (BM) chimeras, B6.SJL-*Ptprc^a^Pepc^b^*/BoyJ (B6.SJL-CD45.1) mice were lethally irradiated (2x 5.0 Gy) and reconstituted with a mixture of 50% B6.SJL-CD45.1 and 50% *Zfp36l1^fl/fl^ Zfp36l2^fl/fl^*(control) or *Zfp36l1^fl/fl^ Zfp36l2^fl/fl^* CD23^cre^ CD45.2 BM cells, corresponding to a total of 3×10^6^ transferred cells. Peripheral blood was sampled to assess reconstitution 8-9 weeks after cell transfer. 1-2 weeks later, chimeric mice were infected with influenza as indicated below.

For immunization, mice received intraperitoneal (i.p.) injection of 200μl sterile PBS containing 100μg of NP(23)KLH adsorbed in 40% v/v Alum (Serva). NP-KLH was emulsified in Alum by rotation for 30 min at room temperature protected from light. The spleen and bone marrow were harvested at day 10 or day 14 post-immunization.

For influenza infection, mice were inoculated intranasally (i.n.) with 10^4^ plaque-forming units of the influenza A/Hong Kong/1/1968/x31 (x31, H3N2) virus strain in 30μL PBS, under inhaled isoflurane anaesthesia using. The mediastinal lymph node and spleen were harvested at day 8 and 15 post-infection.

For *in vivo* BrdU labelling experiments, mice received 2mg BrdU (5-bromo-2′-deoxyuridine, Sigma) i.p. 1.5h before they were culled. For *in vivo* EdU and BrdU double-labelling experiments, mice were first injected i.p. with 1mg EdU (ThermoFisher Scientific). One hour later, the same mice received 2mg BrdU i.p. and were culled 40min later (1h 40min after the first EdU injection).

### B cell isolation and *in vitro* activation assays

Naive B cells were isolated from the peripheral lymph nodes of mice using the B cell isolation kit (Miltenyi Biotec) and cultured in Dutch-modified RPMI-1640 media supplemented with 8% FBS, 1x GlutaMAX (Gibco), 50µM 2-mercaptoethanol, 100units/ml penicillin and 100μg/ml streptomycin (both ThermoFisher Scientific).

For mitogen activation, naive B cells were seeded at 100×10^3^ cells per well in a 96-well plate and cultured with 2.5µg/ml anti-IgM F(ab)2 fragment (Jackson Immune Research), or 2.5µg/ml anti-CD40 antibody (clone: FGK4.5; BioXcell), or a combination thereof, in the presence of 10ng/ml recombinant mouse IL-4 (mIL4; PeproTech).

For inducible germinal centre B cell culture (iGCB), 3T3 cells expressing mouse CD40L and human BAFF (40LB cells^42^) were irradiated (∼120Gγ) and seeded at 100×10^3^ cells per well in a 12-well plate. One day later, 40×10^3^ naive B cells were added to 40LB cells in the presence of 2ng/ml mIL-4. On day 4, B cells were harvested and re-seeded on freshly irradiated 40LB cells in the presence of 5 ng/ml recombinant mouse IL-21 (PeproTech). On day 7, B cells were harvested and used for downstream assays or analysis. 40LB cells were kindly provided by Prof. D. Kitamura, were tested negative for mycoplasma contamination and authenticated for CD40L expression by flow cytometry.

### Flow cytometry

Single cell suspensions from spleens and lymph nodes were prepared by passing the organs through 70μm cell strainers in ice-cold RPMI-1640 with 2% FBS. Single cell suspensions from BM were prepared by flushing the marrow out of the femurs and tibias and passing cells through a 70μm cell strainer in ice-cold RPMI-1640 with 2%FCS. Fc receptors were blocked with monoclonal rat antibody 2.4G2 (BioXcell) and dead cells were excluded with fixable cell viability dye eF780 (ThermoFisher Scientific). For cell surface staining, cells were labelled for 30min at 4°C with the following monoclonal antibodies: anti-CD45.1 (A20), anti-CD45.2 (104), anti-CD19 (1D3 or 6D5), anti-B220 (RA3-6B2), anti-CD4 (RM4-5), anti-CD8 (53-6.7), anti-Ly6C/G (RB6-8C5), anti-CD11c (N418), anti-F4/80 (BM8), anti-IgD (11-26c.2a), anti-IgM (II/41 or RMM-1), anti-IgG1 (A85-1), anti-CD38 (90), anti-GL7 (GL-7), anti-CD95 (Jo2), anti-CD86 (GL-1), anti-CXCR4 (2B11), anti-CD138 (281-2), anti-CD267/TACI (8F10-3). For intracellular staining, cells were fixed with BD Cytofix/Cytoperm for 20 min on ice (BD Biosciences), washed with PBS, frozen at -80°C in 10% DMSO and 90% FBS at least overnight, washed with PBS, re-fixed with BD Cytofix/Cytoperm for 5 min on ice, and washed with BD Perm/Wash buffer (BD Biosciences) prior to permeabilization and intracellular antibody staining. Apoptotic cells were detected using active-caspase 3 antibody (C92-605). NP-specific cells were detected with NP29-PE antibody, or with 4-Hydroxy-3-iodo-5-nitrophenylacetic acid (NIP)15 conjugated to biotin through BSA followed by secondary staining with a streptavidin antibody. EdU was detected with Click-iT Plus AF647 EdU kit (ThermoFisher Scientific), and BrdU with the FITC BrdU kit (BD Biosciences). DNA was stained with 7AAD prior to flow cytometric acquisition.

For replication stress studies, intracellular staining was carried out with the following rabbit monoclonal antibodies: anti-p53-AF647 (1C12 - Cell Signalling), anti-p21-AF647 (F5 - Santa Cruz), or unconjugated rabbit monoclonal anti-pATM (Ser1981, D25E5 - Cell Signalling), anti-cyclin D2 (D52F9 - Cell Signalling), anti-cyclin E2 (E142-Abcam), anti-cyclin B1 (D5C10 - Cell Signalling), anti-pCdc2 (Tyr15; 10A11 - Cell Signalling), anti-CDK2 (78B2 - Cell Signalling), anti-pCHK1 (Ser345; 133D3 - Cell Signalling), or IgG isotype control (Da1E - Cell Signalling); and the following rabbit polyclonal antibodies: anti-pATR (Ser428) and anti-Cdc2 (CDK1; both Cell Signalling). In all cases secondary staining was performed with an AF647-conjugated donkey anti-rabbit IgG(H + L) antibody (Jackson). Data were acquired using a LSR Fortessa Flow Cytometer equipped with 355nm, 405nm, 488nm, 561nm and 640nm lasers (BD Biosciences) and analysed with FlowJo software (TreeStar, version 10.6.1).

### Enzyme-linked immune-spot assay (ELISpot)

For ELISpot, serial dilutions of splenocyte suspensions of immunized mice were added to NP20-BSA-coated or NP2-BSA-coated MultiScreen HA mixed cellulose ester plates (Millipore, Watford, UK) previously washed and blocked with IMDM (Sigma-Aldrich) freshly supplemented with 8% FBS, 2mM GlutaMAX and 50μM 2-mercaptoethanol. Upon overnight incubation, cells secreting anti-NP antibody (ASC) were visualized with HRP-conjugated anti-mouse IgM or IgG1 antibodies (Southern Biotech) followed by AEC staining Kit (Sigma-Aldrich). The numbers of ASCs were quantified using Immunospot S6 Analyzer (Cellular Technology Limited).

### Western blotting

Whole cell lysates were prepared by resuspending B cell pellets in 2x Laemmli buffer containing 5% β2-mercaptoethanol. Protein concentrations were determined by Pierce BCA protein assay (ThermoFisher Scientific). Samples were resolved by 12 % SDS-PAGE and transferred to nitrocellulose membrane using the iBlot2 transfer device (IB21001). Rabbit polyclonal anti-ZFP36L1 (BRF1/2 - Cell Signalling), rat pan-ZFP36 (clone SB1/30.13) and mouse anti-tubulin (DM1A - Sigma) antibodies were detected by incubating membrane with anti-rabbit IgG IRDye680RD (Licor #925-68071), anti-rat IgG IRDye680RD (Licor #925-68076) and anti-mouse IgG IRDye800CM (Licor #926-32210), respectively. Membranes were subsequently scanned with Licor Odyssey CLx using standard methods. Mouse anti-CHK1 (2G1D5 - Cell Signalling) and mouse anti-actin (Sigma) were used in 5% milk prepared in TBS-Tween. Secondary anti-mouse IgG Trueblot HRP-conjugated antibody (eBiosciences) was detected using ECL prime (Cytiva, RPN2232).

### RNA-sequencing

0.5×10^6^ FACS-sorted CD19^+^ B220^hi^ CD138^−^ TACI^−^ iGCB cells (d7) were washed twice with ice-cold PBS and cell pellets were snap frozen in liquid nitrogen. RNA was extracted using the RNeasy Micro Kit (Qiagen) and its quality was assessed on a 2100 Bioanalyser (Agilent). RNA integrity numbers > 8 of total RNA were used to generate cDNA. Opposing strand-specific RNA-sequencing libraries were generated using the NEB PolyA kit (E7490) and the NEB Ultra II stranded kit (E7760) with NEB dual index, according to the manufacturer’s instructions, and sequenced on an Illumina NovaSeq 6000 using 50bp paired-end reads.

The quality of RNA-sequencing data was assessed using FastQC (0.11.9; http://www.bioinformatics.babraham.ac.uk/projects/fastqc/). Reads were trimmed using Trim Galore (0.6.6) and mapped to mouse genome GRCm38 using HiSat2^76^ (2.1.0). Raw counts were calculated using SeqMonk (1.48.0; https://www.bioinformatics.babraham.ac.uk/projects/seqmonk/); this and subsequent analyses were performed using the GRCm38.90 annotation release. DESeq2^77^ (1.39.1) was used to calculate differential RNA abundance between two conditions and performed using default parameters, with “normal” log2 fold change shrinkage.

### Individual-nucleotide resolution UV Cross-Linking and ImmunoPrecipitation (iCLIP)

iCLIP experiments were performed as described in ref.^78^ with lysates of ^mCherry^ZFP36L1 B cells that were cultured for 7 days in iGCB culture condition. WT iGCB cells were used as a negative control. Briefly, iGCB cells were irradiated 1x with ultraviolet (UV) light for cross-linkage of protein-RNA (150 mJ/cm2; Stratalinker 2400). After washing cells in ice-cold PBS, cells were lysed with 50 mM Tris-HCl, pH 7.4, 100 mM NaCl, 1% Igepal CA-630 (Sigma), 0.1% SDS, and 0.5% sodium deoxycholate lysis buffer, and sonicated (3x 15’’). Total protein concentration from extract supernatants was measured by Pierce BCA protein assay and 1.5mg protein were treated for 3 min at 37 °C with 4U Turbo DNAse (Ambion) and 0.4U RNase I (ThermoFisher Scientific). ^mCherry^ZFP36L1-RNA complexes were immunoprecipitated with 20μl RFP-Trap magnetic particles M-270 (Chromotek), which were previously washed 3x in lysis buffer, and rotating overnight at 4°C. Immunoprecipitates were separated by SDS-PAGE. RNA-protein complexes were transferred to a nitrocellulose membrane and isolated after cutting the areas from ∼100 to ∼140 kDa where ^mCherry^ZFP36L1 was expected to be cross-linked to RNAs. After protein digestion, RNA isolation and cDNA library preparation was carried out as described in ref.^78^ using the L3-IR-App adapter to visualize and retrotranscribe the RNA cross-linked to ^mCherry^ZFP36L1. Amplification of iCLIP cDNA libraries was done with 21 PCR cycles. We carried out three biological replicates per condition. Multiplexed iCLIP cDNA libraries were sequenced on a NextSeq500 platform on a 100-bp single-ended mode. iCLIP data were processed using the iCLIP analysis pipeline on the iMaps platform (https://imaps.goodwright.com/ when performed; now hosted on the flow platform: https://app.flow.bio/). Reads were deduplicated based on their barcodes, trimmed using Cutadapt (v2.4)^79^ and mapped to the GRCm38 mouse genome build using STAR (v2.7.0f)^80^. Significant crosslink sites were then determined using iCount. Following iMaps analysis, the gene and feature type to which each cross-link site or cluster was assigned were determined by considering all transcript isoforms from the Ensembl Mus_musculus. GRCm38.90 annotation. If multiple genes or feature types were overlapped, the most likely was chosen by a hierarchy of feature types: 3’UTR > CDS > 5’UTR > intron > non-coding exon > non-coding intron. iCLIP data were visualized using the following application: https://github.com/LouiseMatheson/iCLIP_visualisation_shiny; all cross-links merged across the three replicates were displayed upon filtering for significant sites (FDR < 0.5).

### Transferase-Activated End Ligation sequencing (TrAEL-seq)

TrAEL-Seq experiments were performed using *Zfp36l1^fl/fl^ Zfp36l2^fl/fl^* or *Zfp36l1^fl/fl^ Zfp36l2^fl/fl^*CD23^cre^ B cells that were cultured for 7 days in iGCB culture condition. Agarose-embedded DNA from 2×10^6^ iGCB cells were taken and processed into TrAEL-seq libraries as previously described in ref.^49^, and up-to-date protocols are available from the Houseley lab website (https://www.babraham.ac.uk/our-research/epigenetics/jon-houseley/protocols). TrAEL-seq libraires were sequenced on an Illumina NextSeq500 as high-output 75-bp single end.

For data analysis, UMI deduplicated reads were generated and mapped to the GRCm38 mouse genome as previously described^49^. Detailed information on data processing and scripts are available at https://github.com/FelixKrueger/TrAEL-seq. Mapped reads were analysed through Seqmonk v1.48 (https://www.bioinformatics.babraham.ac.uk/projects/seqmonk/). Minimum mapping quality of 20 was applied, with reads being truncated to 1 bp based on 5’ end and Reads counts were normalised by counts per million reads. Running windows of 80,000 bp and step size of 8,000 bp were used for read density plots. For read polarity plots, running windows of 20,000 bp with step size of 2,000 bp were used and difference quantitation was used calculating (reverse reads / (reverse reads+forward reads)) -50. Active replication initiation zones were determined by the ForkCall program (code found at https://github.com/s-andrews/forkcall) using TrAEL-seq read polarity data, and the plotted read counts were normalised as counts per million reads mapped (RPKM) in 50Kb running windows. Early and late replication initiation zones were retrieved from ref.^51^ (GSE95092).

### Statistics

For mouse experiments, statistical analysis was performed with GraphPad Prism 9.3.1 using two-tailed unpaired Student *t-*test when comparing two groups or one-way ANOVA where indicated. P values < 0.05 were considered statistically significant. Statistics of RNA-sequencing data was conducted in DESeq2 (1.39.1), which uses the Wald test followed by a Benjamini-Hochberg multiple testing correction. For iCLIP data, cross-links were considered significant if adjusted p-value (FDR) was < 0.05. Plots were generated with ggplot2 (3.4.1) and GraphPad Prism 9.3.1.

## Data Availability

RNA-sequencing, TrAEL-Seq and iCLIP data generated in this study are available from the NCBI Gene Expression Omnibus (GEO) repository under the accession code GSE280001. All output files from the iCLIP analysis pipeline are also available on the Flow platform (https://app.flow.bio/projects/361552856316). Flow cytometry data that support the finding of this study are available upon request.

## Author contributions

F.S: conceptualization; methodology; investigation; validation; formal analysis; visualization; data curation; writing - original draft preparation - review & editing; project administration; funding acquisition. A.J.W: methodology; formal analysis; visualization; writing - review & editing. L.S.M: software; formal analysis; visualization; data curation; writing - review & editing. W.S.F, S.E.B, D.J.H: investigation; writing - review & editing. M.S: investigation. M.A.L, J.H: methodology; writing - review & editing. M.T: conceptualization; writing - review & editing; project administration; funding acquisition; supervision.

## Acknowledgments

We thank the Biological Support Unit, Next Generation Sequencing, Flow Cytometry and Bioinformatics facilities of the Babraham Institute, and the Next Generation Sequencing of the CRUK for outstanding technical assistance. We thank Dr. Twm J. Mitchell for his invaluable help with iCLIP experiments. This study was funded by the Biotechnology and Biological Sciences Research Council (BBSRC) (BBS/E/B/000C0427; BBS/E/B/000C0428), the BBSRC Core Capability Grant to the Babraham Institute and a Wellcome Investigator award (200823/Z/16/Z). F.S. was supported by European Molecular Biology Organization (EMBO) Long-Term Fellowship (ALTF 880-2018) and H2020-Marie Skłodowska-Curie Individual Fellowship (841930, B-different).

## Supplementary Figure Legends

**Supplementary Figure 1:**
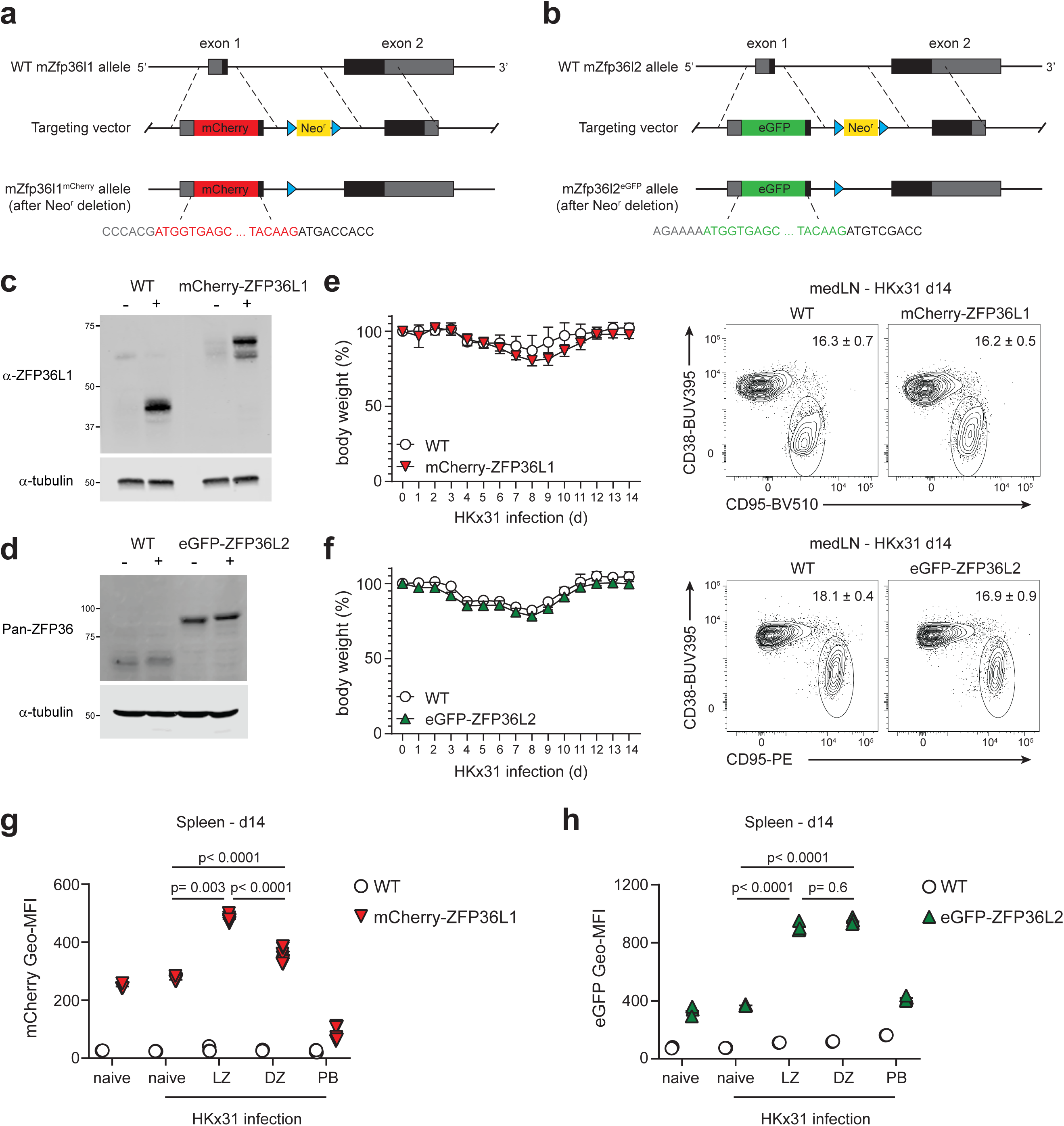
ZFP36L1 and ZFP36L2 fluorescent reporter mice. **a, b)** Schematic illustration of the targeting vector and the resulting engineered mCherry-ZFP36L1 allele **(a)** and eGFP-ZFP36L2 allele **(b)**, where the gene sequences encoding the respective fluorescent reporter proteins have been knocked into the first exons of *Zfp36l1* and *Zfp36l2*, upstream of their ATG start codon. **c, d)** Western blot of ZFP36L1 and mCherry-ZFP36L1**(c)** and ZFP36L2 and eGFP-ZFP36L2 **(d)** expression in B cells that were isolated from homozygous reporter mice or WT control mice. B cells were left untreated (-) or stimulated for 4 hours with 10ng/ml PMA and 1μg/ml ionomycin (+). Equal amounts of protein were loaded per lane. Each blot is a representative of three independent experiments. **e, f)** Mice were i.n. infected with HKx31 influenza virus. Left: body weight loss of initial body weight (100%) in mCherry-ZFP36L1 **(e)**, or eGFP-ZFP36L2 **(f)** mice compared with age and gender matched WT mice. Right: Representative contour plots of GC B cells in medLN 14 days after HKx31 infection. **g, h)** mCherry **(g)** or eGFP **(h)** Geo-MFI was evaluated in naive (CD19^+^ IgD^+^ CD38^+^ CD138^−^) cells, light zone (LZ; CD19^+^ IgD^−^ CD38^−^ CD138^−^ CD95^+^ CXCR4^lo^ CD86^hi^) or dark zone (DZ; CD19^+^ IgD^−^ CD38^−^ CD138^−^ CD95^+^ CXCR4^hi^ CD86^lo^) GC B cells, or plasmablasts (PB; CD19^int/lo^ IgD^−^ CD138^+^) from spleen of mice at day 14 post-infection. Naive B cells of uninfected mice were included as a control. Graphs display mean ±SD of 5-8 mice per group **(e, g)**, or 4-5 mice per group **(f, h)**. Data were pooled from at least two independently performed experiments and were analysed using RM one-way ANOVA with Tukey’s multiple comparison test.

**Supplementary Figure 2:**
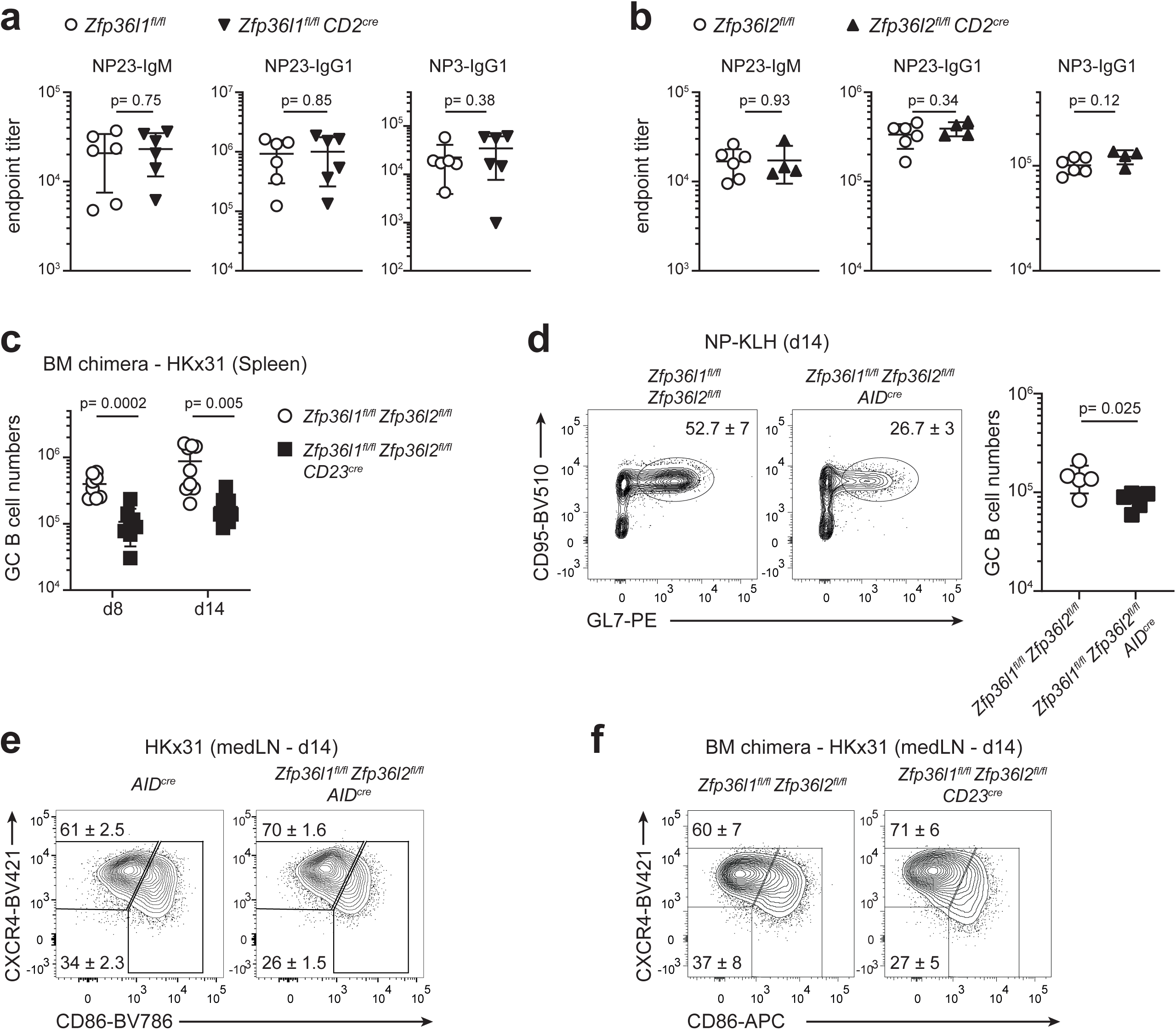
ZFP36L1 and ZFP36L2 are required for GC B cell responses. **a, b)** Serum titres of NP23-IgM, NP23-IgG1 and NP3-IgG1 antibody were analysed by ELISA 14 days after NP-KLH immunization of *Zfp36l1^fl/fl^ CD2^cre^* and *Zfp36l1^fl/fl^* mice **(a)**, or *Zfp36l2^fl/fl^ CD2^cre^* and *Zfp36l2^fl/fl^* mice **(b)**. Unpaired Student’s *t*-test analysis; n= 4-6 mice per group (mean ±SD). Data are representative of at least two independently performed experiments. **(c)** Chimeric mice were infected i.n. with HKx31 influenza virus and spleens were analysed 8 or 14 days later for CD19^+^ IgD^−^ CD38^−^ CD95^+^ GL7^+^ GC B cell numbers. **(d)** Contour plots represent frequency of GC B cells in *Zfp36l1^fl/fl^ Zfp36l2^fl/fl^ AID^cre^* mice and *Zfp36l1^fl/fl^ Zfp36l2^fl/fl^* controls 14 days after NP-KLH immunization. Graph displays mean ±SD of GC B cell numbers. Unpaired Student’s *t*-test analysis; n= 5 mice per group. Data represent two independently performed experiments.

**Supplementary Figure 3:**
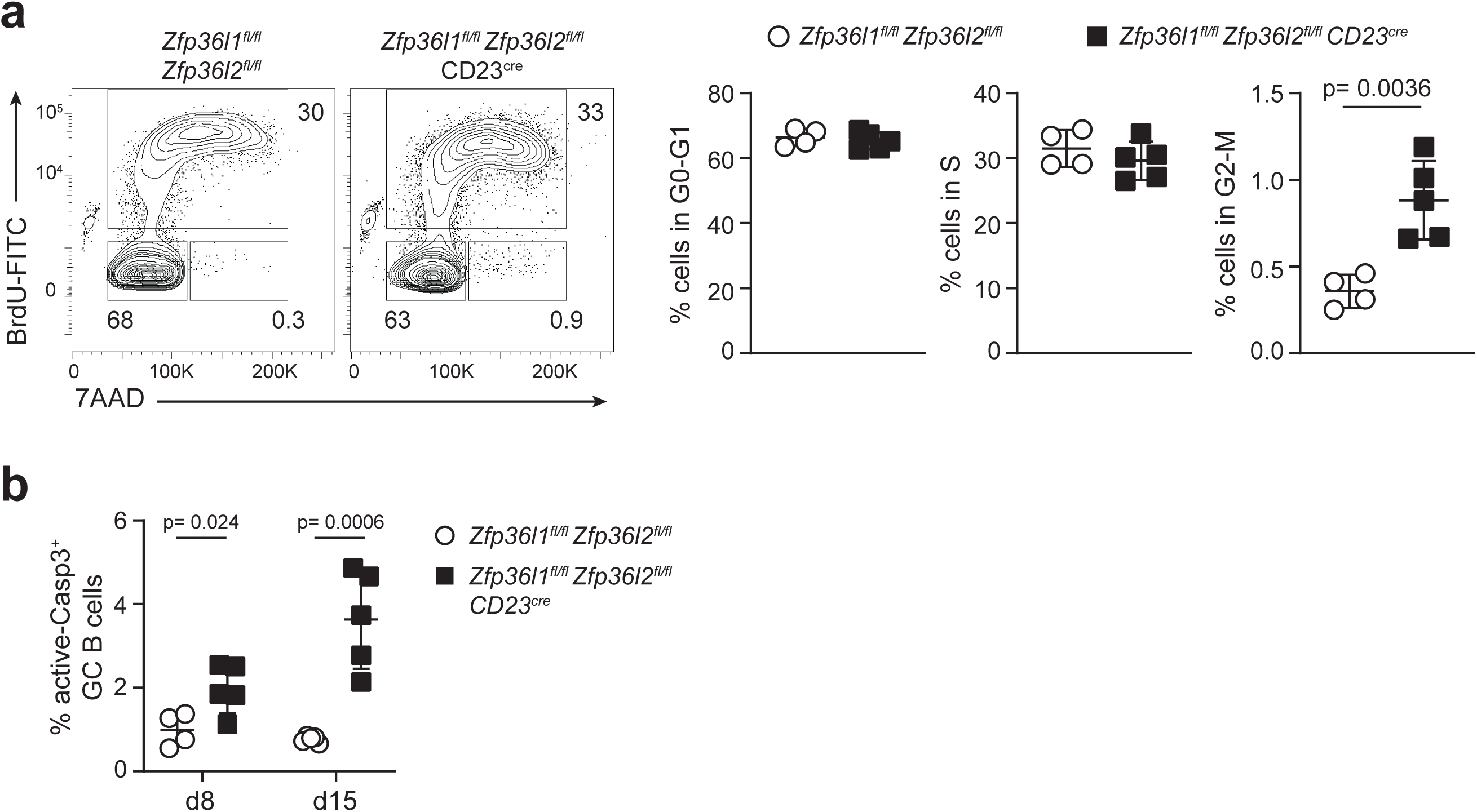
ZFP36L1 and ZFP36L2 control cell cycle progression of GC B cells. **a)** Chimeric mice were analysed 15 days after i.n. HKx31 influenza virus infection. GC B cells (CD19^+^ CD38^−^ CD95^+^ GL7^+^) isolated from medLNs were analysed at various stages of the cell cycle based on flow cytometry staining for BrdU and the membrane-impermeable DNA-intercalating dye 7-AAD. Numbers adjacent to outlined areas indicate frequency of cells in each phase. **b)** Apoptotic GC B cells (gated as in a) in medLNs of chimeric mice that were analysed at day 8 or 15 post-influenza infection.

**Supplementary Figure 4:**
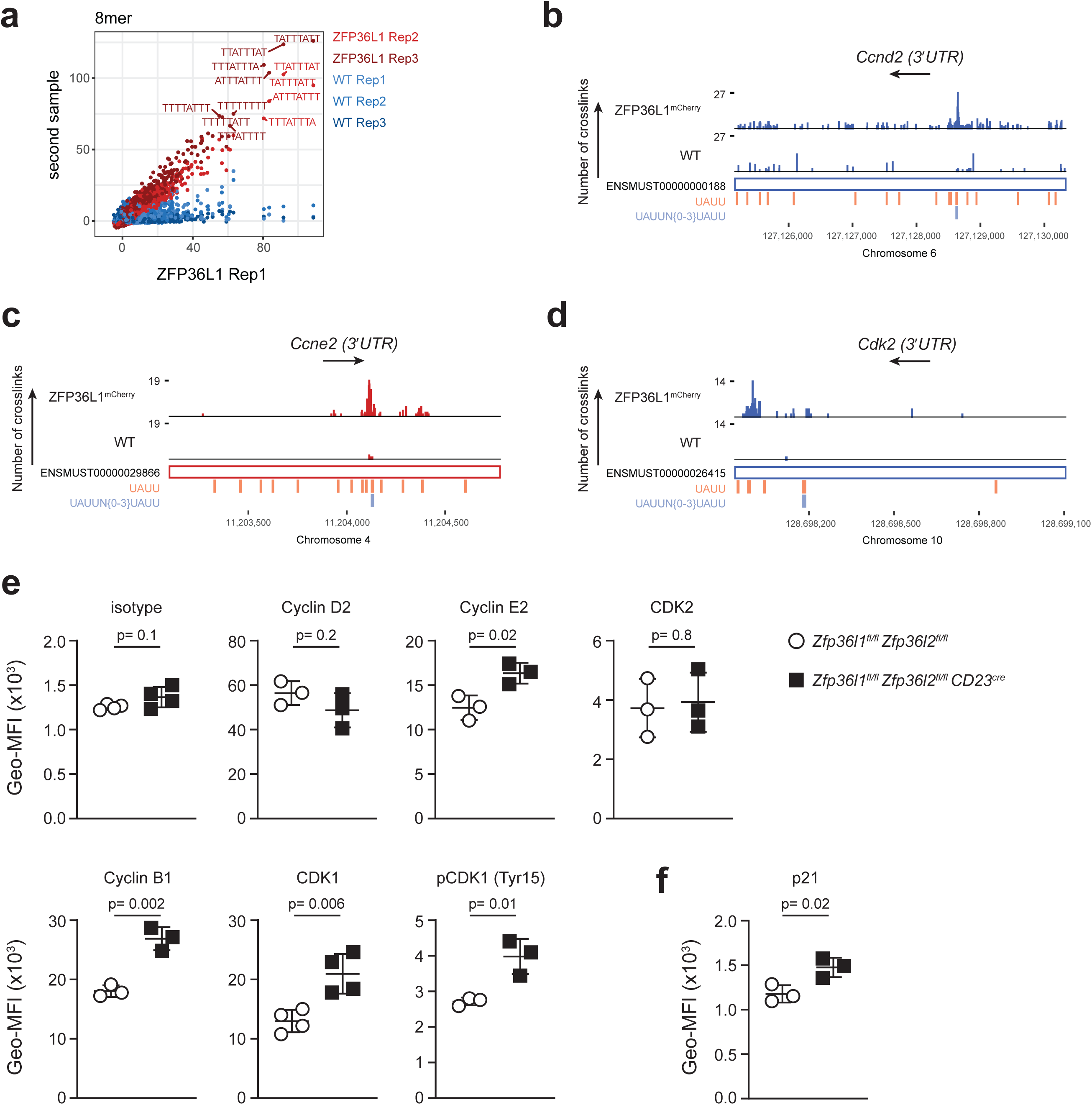
iCLIP identifies ZFP36L1-binding to cell cycle related genes. **a)** Enrichment of 8-nucleotide motifs at sites of mCherry-ZFP36L1binding, identified by iCLIP, relative to randomised sites. Z-scores for replicate 1 of mCherry-ZFP36L1 GC-like B cells are shown on the x-axis; the y-axis represents z-scores for the other replicates of both mCherry-ZFP36L1 and WT GC-like B cells, with each of these comparisons overlaid in a different colour. **b-d)** iCLIP data from GC-like B cells (d7) showing reads across *Ccnd2* **(b)**, *Ccne2* **(c)**, *Cdk2* **(d)** transcripts with an expanded view of their 3’UTR. **e, f)** Graphs depict Geo-MFI of indicated cell-cycle related genes in GC-like B cells. Unpaired Student’s *t*-test analysis; n= 3-4 mice per group (mean ±SD). Data represent two or three independently performed experiments.

**Supplementary Figure 5:**
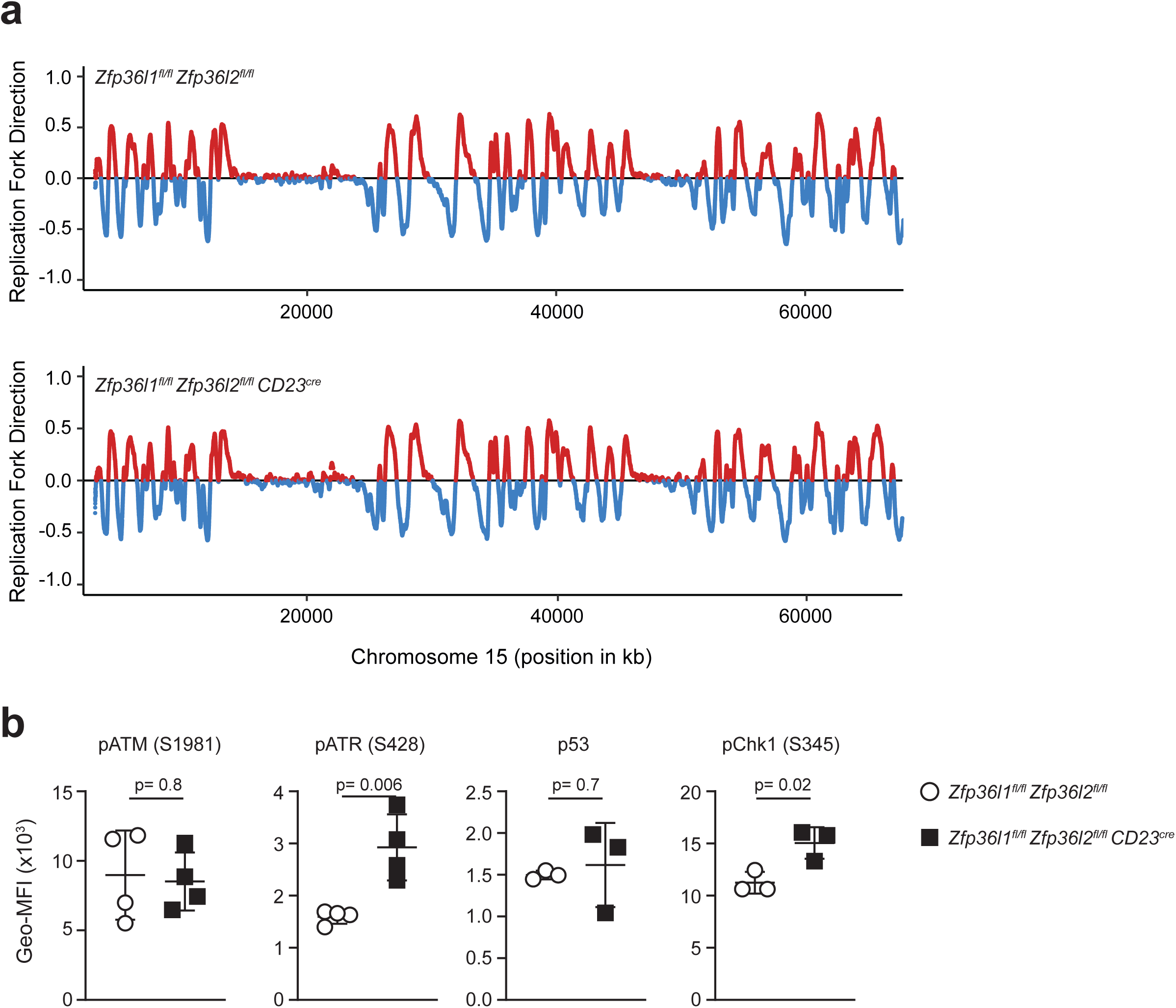
Loss of ZFP36L1 and ZFP36L2 causes replication stress in GC-like B cells. **a)** Read polarity plots for TrAEL-Seq of *Zfp36l1^fl/fl^ Zfp36l2^fl/fl^* control (top) or *Zfp36l1^fl/fl^ Zfp36l2^fl/fl^ CD23^cre^* (bottom) GC-like B cells (d7), across a representative region of the genome (chromosome 15 was randomly selected). Red= forward; blue= reverse. **b)** Graphs depict Geo-MFI of mediators of DNA damage response in *Zfp36l1^fl/fl^ Zfp36l2^fl/fl^* and *Zfp36l1^fl/fl^ Zfp36l2^fl/fl^ CD23^cre^* GC-like B cells. Unpaired Student’s *t*-test analysis; n= 3-4 mice per group (mean ±SD). Data represent two or three independently performed experiments.

## Notes

https://www.ncbi.xyz/geo/query/acc.cgi?acc=GSE280001

### Competing Interest Statement

The authors have declared no competing interest.

